# A compilation of and typology for abundance-, phylogenetic- and functional-based diversity metrics

**DOI:** 10.1101/530782

**Authors:** Samuel M. Scheiner

**Affiliations:** Division of Environmental Biology, National Science Foundation 2415 Eisenhower Ave., Alexandria, VA 22314 USA

## Abstract

Ecologists are faced with an over-abundance of ways to measure biodiversity. In this paper, I provide a compilation of and guide through this ticket of diversity metrics. I present a typology for diversity metrics that encompasses the three commonly considered categories of information: abundance, phylogenetic relationships, and traits (i.e., function). I update and expand previous summaries of diversity metrics. The formulas of those 117 metrics are presented in a standard notation and format that makes it easy to see the mathematical similarities and differences among the metrics. Finally, I propose a standard set of symbols for many of the metrics that makes their properties immediately obvious. This compilation will make it easier for researchers to identify the metric(s) most suited to their needs and will help guide future metric development.

**Disclaimer:** This manuscript is based on work done while serving at (and furloughed from) the U.S. National Science Foundation. The views expressed in this paper do not necessarily reflect those of the National Science Foundation or the United States Government.

## Introduction

Ecologists are faced with an over-abundance of ways to measure biodiversity (Peet 1974, Magurran 1988, Magurran and McGill 2011). Navigating that thicket can be challenging, often resulting in the use of metrics because they are commonly found in the literature, rather than because a given metric is the one most suited to the question being posed. In addition, nearly all commonly used metrics are composites; they consist of more than one component of biodiversity. This composite nature is often hidden by the way the metrics are described or calculated, so determining the correct metric for a given task is not always obvious.

The goals of this paper are fourfold: The first goal is to provide a typology for diversity metrics that encompasses the three commonly considered categories of information: abundancc, phylogenies, and traits (i.e., function). This typology leans heavily on that of Tucker et al. (2017) for phylogenetic diversity metrics, but goes beyond it to further refine their schema and also apply it to trait-based metrics. The second goal is to update the summary of phylogenetic-based diversity metrics of Tucker et al. (2017) and the summary of trait-based diversity metrics of Weiher (2011) as well as proposing new metrics that flow from the new typology. The third goal is to put all of the formulas for these diversity metrics in a standard notation and format that makes it easy to see the mathematical similarities and differences among the metrics. The fourth goal is to propose a standard set of symbols for many of the metrics that makes their properties immediately obvious. The following sections address each of these goals in turn.

## A typology for diversity metrics

Biodiversity consists of a variety of types of information that can be separated and recombined into a myriad of diversity metrics. My proposed typology begins by identifying the basic information components (four types of information, two properties of that information, and three methods for measuring that information), defines a set of elemental diversity metrics from those components, and then shows how those elements can be combined to produce both familiar diversity metrics and new metrics that are analogous to existing metrics using different elements. I start with the four basic types of information: identity, abundance, phylogenetic relationships, and traits. Note that the last item is “traits” rather than “functions” or “functional traits.” While we conventionally speak of “functional diversity,” the “function” part comes from assuming that the traits included in an analysis are the most relevant for the ecological and evolutionary processes that determine the diversity of a community or the effects of those species on ecosystem processes. That assumption is not unwarranted. In general, researchers have an intuitive understanding of which traits are important. That link to processes, however, is almost never actually measured. Rather, we just have the trait measurements. Thus, throughout the rest of this paper I will use the phrase “trait diversity” for the concept that is typically called “functional diversity.”

The four types of information differ in that identity and absolute abundance are aspects of a species that are not mathematically dependent on any other species in an assenblage. In contrast, phylogenetic and trait relationships are aspects that can only be measured relative to other species. They are typically undefined for a monotypic assemblage. Additionally, three of those types of information – abundance, phylogeny, traits – evince two properties: magnitude and variability that each can vary independently and which are labeled as follows (Scheiner et al. 2017a):

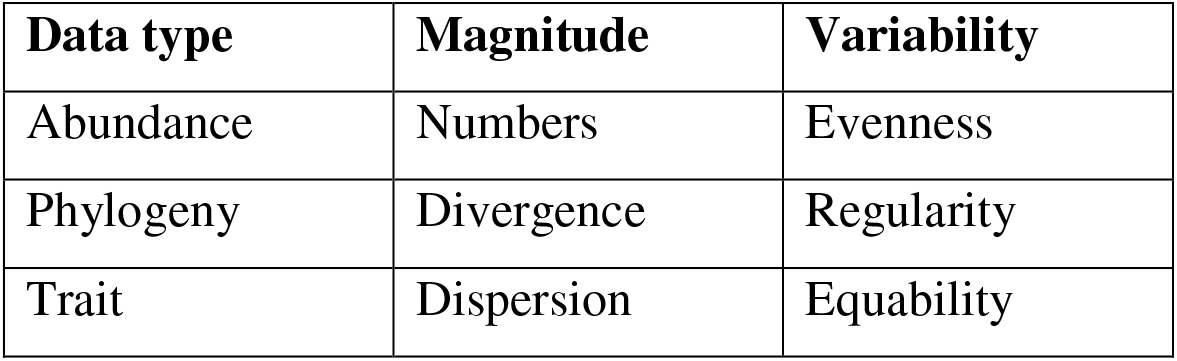

Magnitude is how much each of the species in an assemblage manifests some property. For abundance, magnitude (numbers) is typically the total number of individuals of a particular species, although it can be measured in a variety of other ways such as frequency of occurrence, biomass, or geographic range. For phylogeny, magnitude (divergence) is the amount of evolutionary differentiation of a particular species from other species (Figure 1A). For traits, magnitude (dispersion) is the amount of difference in trait values of a particular species from other species (Figure 1B). Variability quantifies the extent to which magnitudes differ among those species. For abundance, variability (evenness) is the similarity in the (relative) number of individuals of each species. For phylogeny, variability (regularity) is the extent to which species are equally divergent. For traits, variability (equability) is the extent to which species are equally different from each other in trait values.

**Figure 1.**
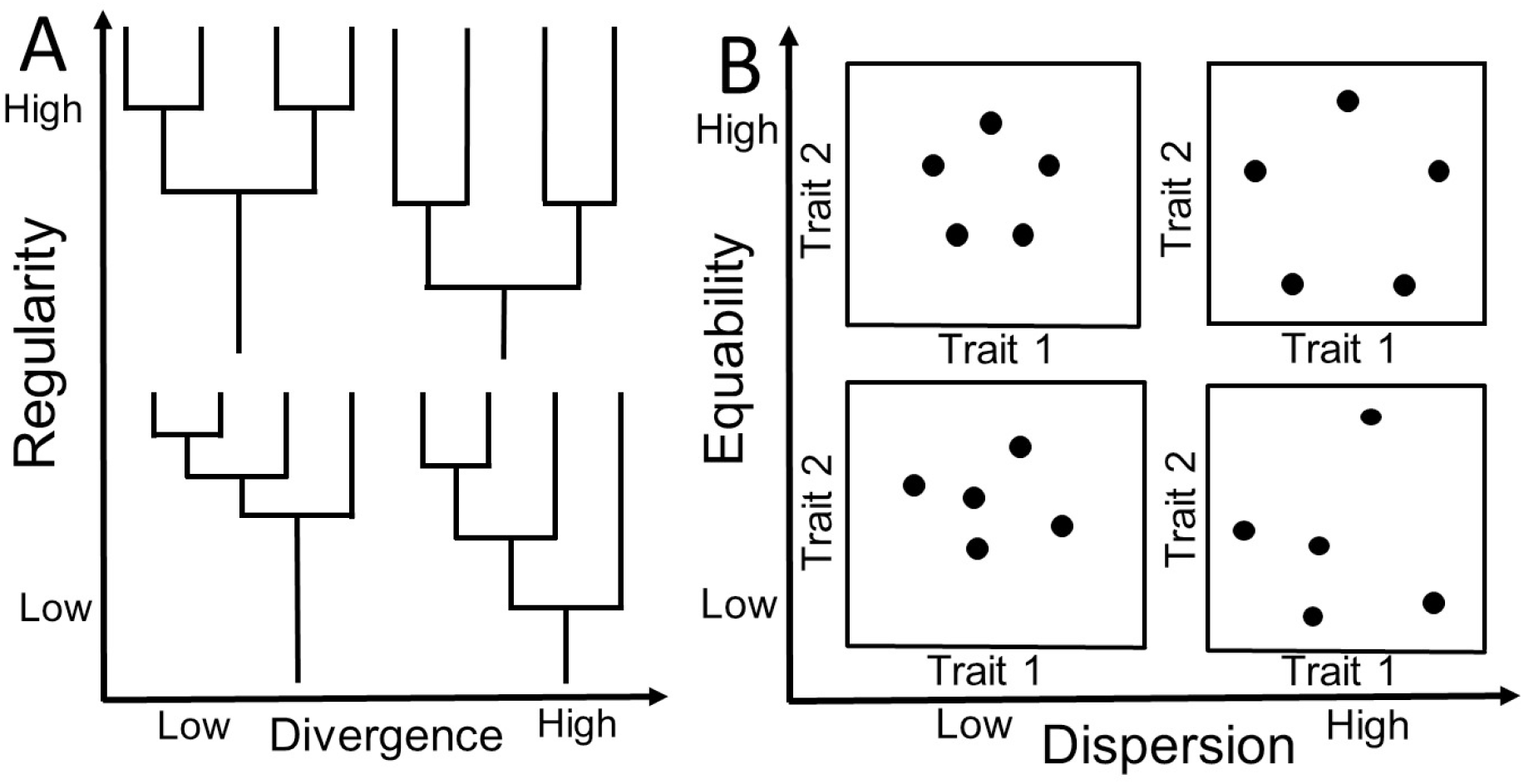
(A) Four phylogenies showing how magnitude (divergence) and variability (regularity) can vary independently. A phylogeny with higher divergences has longer branch lengths towards the tips. A phylogeny with higher regularity has more symmetrical or similar length branches. (B) Four assemblages showing how magnitude (dispersion) and variability (equability) can vary independently. Species with higher dispersions have greater distances among the species in trait space. Species with higher equability have more similar distances among the species.

Because both phylogenetic and trait information are determined relative to other species, there are three methods for measuring these properties: total, pairwise, and nearest-neighbor. For phylogenetic information, total metrics are based on mseasures of the total branch lengths from root to tip for each species. Most commonly, shared branch lengths are divided by what are called “fair proportions” (Isaac et al. 2007) although other divisions are possible (Cadotte and Jonathan Davies 2010). Both pairwise and nearest-neighbor metrics are based on the lengths of branches from one tip of the phylogeny to another, with the former comprising all pairwise lengths and the latter just the smallest length. They both differ from total metrics in that they do not include any shared branches. For example, in Figure 1A, the single root branch in each of the phylogenies would not be included in any pairwise or nearest-neighbor metrics. Thus, for total metrics the sum of the values for each species always equal the total branch length; that equivalence does not hold for pairwise or nearest-neighbor metrics.

For trait information, total metrics are based on the mean distance of each species from all others, pairwise metrics are based on the individual pairs of distances between each species, and nearest-neighbor metrics are based on just the smallest distance for each species. (Analogous quantities can be computed if the traits are categorical, rather than continuous.) Both total and pairwise metrics have the same sums and overall means so that magnitude metrics do not differ; they do differ for variability metrics. Trait diversity metrics can also be separated based on two concepts: uniqueness and combinatorics (see Figure 1 of Scheiner et al. 2017b). For the uniqueness concept, if data are categorical, trait diversity is greatest when each species in an assemblage has a unique set of trait attributes; if the data are continuous, diversity is greatest when the species are as far apart from each other as possible. For the combinatorics concept, if data are categorical, diversity is greatest when an assemblage contains species that have every possible combination of trait attributes; if the data are continuous, diversity is greatest when dispersion is as compact as possible while equalizing the minimum distances between species.

Leinster and Cobbold (2012) provide a list of preferred properties for diversity metrics based on consideration of effective numbers, modularity, replication, symmetry, the effect of absent species, the effect of identical species, monotonicity, how metrics simplify when abundance, phylogeny, or trait values are not included (naive model), and the range of the metric. Some of those properties (e.g., effective numbers, data range) hold only for metrics that combine species richness with other elements. So not all diversity metrics satisfy their criteria. However, their criteria are useful for examining the properties of various metrics. The list of metrics presented in this paper is meant to be comprehensive rather than prescriptive. I take no position on whether all of the properties of Leinster and Cobbold are required for any given metric.

## A summary of diversity metrics

I begin my summary of diversity metrics by defining 14 basic elements, the 13 cells in Table 1 plus identity information. The most common type of identity is “species” but other units are possible (e.g., genus, genotype). When the units are species, then identity diversity equals species richness (S). Those 14 basic elements can be measured in more than one way (e.g., the three different metrics for equability based on nearest-neighbor distances). After combining those elements in various ways, the result is 117 different metrics. That number does not include all metrics already in the literature, although it does include the commonly used ones, as well as many new metrics.

At this point, you may well ask yourself: “Does ecology really need this many ways to measure biodiversity?” My purpose here is not to overwhelm the reader. Quite the opposite. My purpose is to show that all of these metrics are simply combinations of the 14 basic elements, and even those basic elements reduce to just three categories (four types of information, two properties, and three measurement methods).

**Table 1.** Metrics of thirteen of the basic elements (plus identity). Metrics in this and subsequent tables in **bold** are newly proposed; metrics in *italics* are implied by previously proposed metrics. See Table A1 for the formulas for each metric, along with the name, alternative symbols, the quantities that each measure, and the source.

|  | Magnitude |  |  | Variability |  |  |
| --- | --- | --- | --- | --- | --- | --- |
| Data type | Total | Pairwise | Nearest | Total | Pairwise | Nearest |
| Abundance | $M(A)$ | --- | --- | ${}^qE(A)$ | --- | --- |
| Phylogeny | $M(P_T)$<br>$AvPD$ | $M(P_P)$<br>$J$ PSV | $M(P_N)$ | ${}^qE(P_T)$<br>**$C(P_T)$** | **${}^qE(P_P)$**<br>**$C(P_P)$** | **${}^qE(P_N)$**<br>**$C(P_N)$** |
| Trait | $M(T_{T/P})$ | | $M(T_N)$ | ${}^qE(T_T)$ | ${}^qE(T_P)$ | ${}^qE(T_N)$<br>**$C(T_N)$**<br>FNEve |

Organizing the metrics in this fashion accomplishes three goals. First, it identifies which metrics are based on the same types of information, properties, or measurement methods. If you wish to compare phylogenetic and trait diversity for a set of species, it does not make sense to compare a measure of pairwise divergence with a measure of nearest-neigbor equability. The tables below make it easy to identify comparable metrics.

Second, it helps distinguish between different types of combinations of elements, for example, a measure of abundance-weighted phylogenetic diversity [^q^DA(P_T_), which combines elements of identity, numbers, and total regularity] as compared to a measure of phylogenetic-weighted abundance diversity [^q^DP_T_(A), which combines elements of identity, evenness, and total divergence] (see table A5). Recognizing that these two metrics combine different basic elements prevents confusion or inappropriate comparisons, especially across studies done by different researchers.

Third, it identifies “missing” metrics. Imagine Tables 1–5 without any of the metrics in italic or bold font. That is how I began this compilation. The metrics in italic are ones that are implied by previously published metrics. The most obvious are measures of regularity or equability that are derived from Hill-based diversity metrics by dividiing by S (e.g., ^q^E(T_T_), Table 1). The metrics in bold are ones that were suggested by analogy with existing metrics. For example, metrics had been proposed for divergence based on total and pairwise measures and for regularity based on total measures (Table 2). Those measures suggested the possibility of additional metrics of nearest-neighbor divergence, pairwise regularity, and nearest-neighbor regularity based on simple substitutions or modifications of existing formulas. Once I began this process, it became straightforward to continue until all of the cells in Tables 1–5 contained at least one metric. Even if a cell already contained a metric, others were added if they were based on combinations of basic elements that were not previously considered, or if based on different concepts of magnitude or variability. For the most part, the metrics are as originally presented. My only systematic deviation from that practice is that all of the variance-based metrics of Tucker et al. (2017) were converted to standard deviations so as to make units comparable between magnitude and variability (Table A3).

**Table 2.** Metrics that combine species richness with one other basic element (see Table A2 for the formulas).

|  | Magnitude |  |  | Variability |  |  |
| --- | --- | --- | --- | --- | --- | --- |
| Data type | Total | Pairwise | Nearest | Total | Pairwise | Nearest |
| Abundance | $N$ | --- | --- | ${}^qD(A)$ | --- | --- |
| Phylogeny | $\Sigma(P_T)$<br>$M(PR)$ | $\Sigma(P_P)$<br>PSR | **$\Sigma(P_N)$** | ${}^qD(P_T)$ | **${}^qD(P_P)$** | **${}^qD(P_N)$** |
| Trait | $\Sigma(T_{T/P})$<br>FRic | $\Sigma(T_{T/P})$ | **$\Sigma(T_N)$**<br>FD | ${}^qD(T_T)$ | ${}^qD(T_P)$ | ${}^qD(T_N)$ |

**Table 3.** Metrics that combine magnitude and variability of the same data type (see Table A3 for the formulas).

| Data type | Total | Pairwise | Nearest |
| --- | --- | --- | --- |
| Phylogeny | $\sigma(P_T)$ | $\sigma(P_P)$<br>$\sigma A(P_P)$ | $\sigma(P_N)$<br>$\sigma A(P_N)$ |
| Trait | **$\sigma(T_T)$** | ${}^qDM(T_P)$<br>${}^qDMA(T_P)$ | $\sigma(T_N)$ |

**Table 4.** Metrics that combine abundance with one other basic element involving phylogeny or traits (see Table A4 for the formulas).

|  |  | Magnitude |  |  | Variability |  |  |
| --- | --- | --- | --- | --- | --- | --- | --- |
| Data types |  | Total | Pairwise | Nearest | Total | Pairwise | Nearest |
| Phylogeny | Abundance magnitude | $\mathbf{MA(P_T)}$<br>$\text{avPD}_{\text{Ab}}$ | $\mathbf{MA(P_P)}$ | $\text{MA}(P_N)$ | $\text{MP}_T(\mathbf{A})$<br>${}^q\text{E}_I\mathbf{A(P_T)}$<br>${}^q\mathbf{EA(P_T)}$ | ${}^q\mathbf{EA(P_P)}$ | ${}^q\mathbf{EA(P_N)}$<br>PEve |
| | Abundance variation | ${}^q\mathbf{EP_T(A)}$ | ${}^q\mathbf{EP_P(A)}$ | ${}^q\mathbf{EP_N(A)}$ | ${}^qE(AP_T)$ | ${}^q\mathbf{E_I(AP_P)}$<br>${}^q\mathbf{E(AP_P)}$ | ${}^q\mathbf{E(AP_N)}$ |
| Trait | Abundance magnitude | $\mathbf{MA(T_T)}$<br>$\text{M}_I(\mathbf{T_T})$<br>FDis | $\mathbf{MA(T_P)}$ | $\text{MA}(T_N)$ | $\mathbf{MT_T(A)}$<br>${}^q\mathbf{EA(T_T)}$<br>FDiv | ${}^q\mathbf{EA(T_P)}$ | ${}^q\mathbf{EA(T_N)}$<br>FEve |
| | Abundance variation | ${}^q\mathbf{ET_T(A)}$ | ${}^q\text{ET}_P(\mathbf{A})$<br>${}^qFE(Q)$ | ${}^q\mathbf{ET_N(A)}$ | ${}^q\mathbf{E(AT_T)}$<br>${}^q\mathbf{E(AP_TT_T)}$ | ${}^q\text{E}_I(\mathbf{AT_P})$<br>${}^q\mathbf{E(AT_P)}$ | ${}^qE(AT_N)$<br>${}^qE(AP_TT_N)$ |

**Table 5.** Metrics that combine species richness and abundance with one or more other basic elements involving phylogeny or traits (see Table A5 for the formulas).

|  |  | Magnitude |  |  | Variability |  |  |
| --- | --- | --- | --- | --- | --- | --- | --- |
| Data types |  | Total | Pairwise | Nearest | Total | Pairwise | Nearest |
| Phylogeny | Abundance magnitude | $\Sigma(\mathbf{AP_T})$<br>$\Sigma\mathbf{A(P_T)}$ | $\Sigma(\mathbf{AP_P})$<br>PSE | $\Sigma\mathbf{A(P_N)}$ | ${}^q\text{D}_I\mathbf{A(P_T)}$<br>${}^q\mathbf{DA(P_T)}$ | ${}^q\mathbf{DA(P_P)}$ | ${}^q\mathbf{DA(P_N)}$ |
| | Abundance variation | ${}^q\text{DP}_T(\mathbf{A})$ | ${}^q\text{DP}_P(\mathbf{A})$ | ${}^q\mathbf{DP_N(A)}$ | ${}^q\text{D}(\mathbf{AP_T})$ | ${}^q\mathbf{D_I(AP_P)}$<br>${}^q\mathbf{D(AP_P)}$ | ${}^q\mathbf{D(AP_N)}$ |
| Trait | Abundance magnitude | $\Sigma\mathbf{A(T_T)}$ | $\Sigma\mathbf{A(T_P)}$ | $\Sigma\mathbf{A(T_N)}$ | ${}^q\mathbf{DA(T_T)}$ | ${}^q\mathbf{DA(T_P)}$ | ${}^q\mathbf{DA(T_N)}$ |
| | Abundance variation | ${}^q\mathbf{DT_T(A)}$ | ${}^q\text{DT}_P(\mathbf{A})$ | ${}^q\mathbf{DT_N(A)}$ | ${}^q\mathbf{D(AT_T)}$<br>${}^q\mathbf{D(AP_TT_T)}$ | ${}^q\text{D}_I(\mathbf{AT_P})$<br>${}^q\mathbf{D(AT_P)}$ | ${}^q\mathbf{D(AT_N)}$<br>${}^q\mathbf{D(AP_TT_N)}$ |

Fourth, my typology makes it easier for researchers to develop new metrics if none of the existing metrics are best suited for the task at hand. Additional metric might be implied by analogy with those presented here. Or, the basic elements might be combined in yet new ways. Or different types of basic elements might be developed, for example other ways of measuring variability.

The effective number of species are the number of species that an assemblage would contain if all species had equal abundances, divergences, or dispersions (Jost 2006). They are obtained by multiplying a measure of evenness, regularity, or equability by species richness (Jost 2010). The metrics in Tables 2 and 5 are the only ones that might represent effective numbers. Not all of those metrics result in measures of the effective number of species, however. Only if the other element(s) of the metric have a range of (0,1] will the resulting combination have a range of (0,S]. So species richness (identity) is a necessary, but not sufficient, ingredient for a measure of effective numbers.

## Standardizing diversity metric formulas

All of the formulas in the appendix use a consistent set of symbols for all parameters. Doing so avoids creating confusion by the use of the same symbol for different concepts (e.g., using “*d*” for both branch length and trait-space distance). For the most part, the symbols used are those typically found in previous publications, except where the same symbol was being used for differen concepts. Where possible, subscripts are used to distinguish between related parameters (e.g., *min* for metrics involving nearest-neighbor divergences or distances). The formulas are also presented in as atomistic a fashion possible (e.g., denominators show full summations rather than a composite sum). Doing so makes it easy to see the components of each formula, avoids confusion, and limits the number of symbols needed. I urge other researchers to further this usage to make as clear as possible the relationship of any new metric to current metrics.

## A proposed set of symbols for diversity metrics

I am proposing a consistent symbology for many of the diversity metrics. This system is designed to convey information about the content of the metric; that is, the symbol is more than just an arbitrary designation. The system is applied primarily to metrics that are based on Hill diversity (Hill 1973), but also those that rely on standard mathematical operations (averages, standard deviations, and coefficients of variation). I developed this system in assembling the metrics in this paper. Especially as new metrics were being described, I discovered that previous symbols, including ones in my own papers, were inadequate for creating a unique symbol for each metric. In addition, for some metrics there was no single, standard symbol. The proposed system conveys the three categories of information as follows:

1. Type of information: A = abundance, P = phylogenetics, T = traits
2. Property as determined by the mathematical function:

Magnitude: M = mean, Σ = sum
Variability: σ = standard deviation, C = coefficient of variation, D = Hill diversity, E = Hill evenness
For Hill functions, the superscript q = exponent (typically 0, 1, or 2)
For trait information, the subscript I = operation performed on individuals within species
3. Type of measurement for phylogenetic or trait information denoted as a subscript: T = total, P = pairwise, N = nearest-neighbor

If the symbol for a type of information is inside the parenthesis of a function, the function is performed on that parameter. If the symbol is outside the parenthesis, that parameter is a weighting factor. For example, ^q^DT_P_(A) is Hill diversity of abundance information weighted by pairwise trait information; ^q^DA(T_P_) is Hill diversity of pairwise trait information weighted by abundance information; ^q^D(AT_P_) is Hill diversity of both abundance and pairwise trait information. Not included here, because I do not list their formulas, are symbols designating ecological hierarchies (α-. β-, and γ-diversity). Those would be added as subscripts of the mathematical function.

I urge the use of a consistent set of symbols to make it easier for the reader to understand what metric is being used, especially if the formula for that metric is not given in the publication. I am well aware that getting authors to do so is difficult. In one case, one of my own metrics was used in a paper with a notation that differed from the one in my original publication. As my publication listed several different metrics, it was not obvious which one was being used. Editors and reviewers are critical for encouraging usage consistency.

## Concluding remarks

I wish to be clear that in this paper I am not advocating the use of any specific metrics. Rather, my goal is to be as comprehensive as possible so that you can identify the metric(s) that are most appropriate for the question that you wish to address. This task is especially important for metrics that are composites of two or more elements, as it is not always obvious what elements are being combined. For example, is this metric measuring abundance diversity weighted by phylogenetic information, or phylogenetic diversity weighted by abundance information? Such identification ensures that comparisons among diversities measured with different types of information (e.g., phylogenetic and trait information) are done with metrics based on similar properties (e.g., both are based on nearest-neighbor data). Even more important, it points out that certain types of comparison are not informative. A correlation of Hill diversity metrics of phylogenetic and trait data is such a comparison. Both metrics include species richness as a component so that any correlation is (partially) based on a correlation of a parameter with itself. Instead, the correlation should be done using Hill evenness metrics, which do not contain overlapping elements.

My compilation is incomplete in several regards. First, I have undoubtably left out some metrics despite my attempts to ferret them out. Second, there are other aspects of biodiversity that are not addressed here; my focus is exclusively on metrics relevant to species within communities. Nor does the compilation consider other types of measures, such as entropy (Jost 2006). Third, all of the formulas presented here assume no sampling bias. Bias-corrected formulas for some of the metrics can be found in (Colwell and Coddington 1994, Chao and Jost 2012, Chao et al. 2013, Chao et al. 2015) among other publications. Fourth, I do not deal with hierarchical data structures and metrics of α-diversity (mean subsample) and β-diversity (among subsample). Again, formulas for hierarchical data structures for some of the metrics can be found in various other publications (e.g., Tuomisto 2010, Chao et al. 2012, Chiu et al. 2014, Pavoine et al. 2016, Scheiner et al. 2017b, Tucker et al. 2017, Podani et al. 2018).

## Appendix

In all tables, previously used symbols are given in brackets. The source given is the first occurrence of a metric in the literature, even when that metric was subsequently, independently derived. Symbols use in the tables are as follows:

S: the number of species
*n_i_*: the number of individuals of species *i*
*N*: the total number of individuals in an assemblage
*T*: the time-depth of a cladogram
*B*: the number of branches on a cladogram
*N_b_*: the total number of individuals of all species that share the *b*th branch segment on a cladogram
*L_b_*: the length of the bth branch segment on a cladogram
*L_ib_*: the proportional share of the *b*th branch segment of species *i*
*L_i_*: the total proportional branch length share of species *i*
*φ_ij_*: the total branch length between species *i* and *j*
*φ_i min_*: the shortest branch length between species *i* and all other species
*ψ_ij_*: the amount of unshared branch length between species *i* and *j*, = 1 − [*c_ij_*/√(*c_ii_ c_jj_*)], where *c_ii_*(*c_jj_*) = the sum of the branch lengths for species *i*(*j*), and *c_ij_* is the sum of the branch lengths from the root to the most recent common ancestor of species *i* and *j*. [Note that *ψ_ij_* = *φ_ij_* for the alternative definition, = 0.5(*c_ii_* + *c_jj_* − *c_ij_*).]
*d_i_*: the mean standardized trait distance of species *i* and all other species
*d_ij_*: the standardized trait-space distance between species *i* and *j*
*d_i min_*: the smallest standardized trait-space distance between species *i* and all other species
*d_kl_*: the standardized trait-space distance between individuals *k* and *l* ignoring species identity
*δ_i_*: the distance of species *i* from the abundance-weighted centroid of the species trait-space distribution
Δ*d*: the sum of abundance-weighted deviances from the centroid of the trait-space convex hull
Δ|*d*|: the sum of absolute abundance-weighted deviances from the centroid of the trait-space convex hull
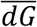: the mean distance from the centroid of the trait-space convex hull
*μ_i min_*: the length of the *l*th branch connecting species *i* and *j* of a minimum spanning tree constructed from phylogenetic or trait distances
q: the exponent of the Hill function

**Table A1.** Metrics of fourteen of basic elements

| Metric | Symbol | Description | Formula | Quantities | Source |
| --- | --- | --- | --- | --- | --- |
| **A. Identity** |  |  |  |  |  |
| Species richness | S | Number of taxonomic units | NA | Identity | NA |
| **B. Magnitude** |  |  |  |  |  |
| Mean abundance | M(A) | Mean number of individuals | $\frac{\sum_{i=1}^S n_i}{S}$ | Numbers | NA |
| Mean proportional divergence | M(P<sub>T</sub>)<br>[M(P)] | Time-depth standardized mean species divergence | $\frac{\sum_i^S \sum_{b(S_i)} L_{ib}}{T \times S}$ | Total divergence | (Scheiner et al. 2017a) |
| Average phylogenetic diversity | AvPD | Mean species divergence | $\frac{\sum_i^S \sum_{b(S_i)} L_{ib}}{S}$ | Total divergence | (Clarke and Warwick 2001) |
| Mean pairwise divergence | M(P<sub>P</sub>)<br>[MPD<br>AvTD Δ<sup>+</sup>] | Mean phylogenetic divergence | $\frac{\sum_{i=1}^S \sum_{j \neq i}^S \varphi_{ij}}{S(S-1)}$ | Pairwise divergence | (Clarke and Warwick 1998) |
| Intensive quadratic entropy | J | Mean phylogenetic divergence | $\frac{\sum_{i=1}^S \sum_{j=1}^S \varphi_{ij}}{S^2}$ | Pairwise divergence | (Izsák and Papp 2000) |
| Phylogenetic species variability | PSV | Mean amount of unshared branch length | $\frac{\sum_{i=1}^S \sum_{j \neq i}^S \psi_{ij}}{S(S-1)}$ | Pairwise divergence | (Helmus et al. 2007) |
| Mean nearest taxon distance | M(P<sub>N</sub>)<br>[MNTD] | Mean shortest phylogenetic distance between taxa | $\frac{\sum_{i=1}^S \varphi_{i \min}}{S}$ | Nearest divergence | (Webb et al. 2002) |
| Mean trait dispersion | M(T<sub>T/P</sub>)<br>[MFAD] | Mean pairwise trait distance | $\frac{\sum_{i=1}^S \sum_{j=1}^S d_{ij}}{S(S-1)}$ | Total (or pairwise) dispersion | (Scheiner et al. 2017b) |
| Mean nearest trait dispersion | M(T<sub>N</sub>)<br>(meanNND) | Mean shortest trait distance between taxa | $\frac{\sum_{i=1}^S d_{i \min}}{S}$ | Nearest dispersion | (Ricklefs and Travis 1980) |
| **C. Variability** |  |  |  |  |  |
| Hill evenness | <sup>q</sup>E(A) | Equality of the number of individuals | $\left( \sum_{i=1}^S \left( \frac{n_i}{\sum_{i=1}^S n_i} \right)^q \right)^{1/(1-q)} / S$ | Evenness | (Hill 1973) |
| Hill regularity | <sup>q</sup>E(P<sub>T</sub>) | Symmetry of branch lengths | $\left( \sum_{i=1}^S \left( \frac{\sum_{b(S_i)} L_{ib}}{\sum_i^S \sum_{b(S_i)} L_{ib}} \right)^q \right)^{1/(1-q)} / S$ | Total regularity | (Implied by Scheiner 2012) |
| CV of regularity | $C(P_T)$ | Coefficient of variation of branch lengths | $\sqrt{\left(\sum_{i=1}^S \left(\sum_{b \in b(S_i)} L_{ib} - \frac{\sum_{b \in b(S_i)} L_{ib}}{S}\right)^2\right) / (S-1)} / \frac{\sum_{i=1}^S \sum_{b \in b(S_i)} L}{S}$ | Total regularity | This paper |
| Hill pairwise regularity | ${}^qE(P_P)$ | Evenness in pairwise phylogenetic divergence | $\left(1 + \sqrt{1 + 4 \left(\sum_{i=1}^S \sum_{j=1}^S \left(\frac{\varphi_{ij}}{\sum_{i=1}^S \sum_{j=1}^S \varphi_{ij}}\right)^q\right)^{1/(1-q)}}\right) / 2S$ | Pairwise regularity | This paper |
| CV of pairwise distances | $C(P_P)$ | Coefficient of variation of pairwise phylogenetic divergence | $\sqrt{\sum_{i=1}^S \sum_{j \neq i}^S \left(\varphi_{ij} - \frac{\sum_{i=1}^S \sum_{j \neq i}^S \varphi_{ij}}{S(S-1)}\right)^2 / S(S-1)} / \frac{\sum_{i=1}^S \sum_{j \neq i}^S \varphi_{ij}}{S(S-1)}$ | Pairwise regularity | This paper |
| Nearest taxon Hill regularity | ${}^qE(P_N)$ | Equality of the nearest pairwise phylogenetic divergence | $\left(\sum_{i=1}^S \left(\frac{\varphi_{i \min}}{\sum_{i=1}^S \varphi_{i \min}}\right)^q\right)^{1/(1-q)} / S$ | Nearest regularity | This paper |
| CV of nearest taxon distances | $C(P_N)$ | Coefficient of variation of nearest pairwise phylogenetic divergence | $\sqrt{\sum_{i=1}^S \left(\varphi_{i \min} - \frac{\sum_{i=1}^S \varphi_{i \min}}{S}\right)^2 / S} / \frac{\sum_{i=1}^S \varphi_{i \min}}{S}$ | Nearest regularity | This paper |
| Hill equability | ${}^qE(T_T)$ | Equality in mean trait distance | $\left(\sum_{i=1}^S \left(\frac{d_i}{\sum_{i=1}^S d_i}\right)^q\right)^{1/(1-q)} / S$ | Total equability | (Implied by Presley et al. 2014) |
| Hill pairwise equability | ${}^qE(T_P)$<br>[${}^qE(T)$] | Equality in pairwise distances | $\left(1 + \sqrt{1 + 4 \left(\sum_{i=1}^S \sum_{j=1}^S \left(\frac{d_{ij}}{\sum_{i=1}^S \sum_{j=1}^S d_{ij}}\right)^q\right)^{1/(1-q)}}\right) / 2S$ | Pairwise equability | (Scheiner et al. 2017b) |
| Hill nearest equability | ${}^qE(T_N)$ | Equality in nearest distances | $\left(\sum_{i=1}^S \left(\frac{d_{i \min}}{\sum_{i=1}^S d_{i \min}}\right)^q\right)^{1/(1-q)} / S$ | Nearest equability | (Implied by Scheiner 2012) |
| CV of nearest neighbor distance | $C(T_N)$ | Coefficient of variation in nearest distances | $\sqrt{\sum_{i=1}^S \left(d_{i \min} - \frac{\sum_{i=1}^S d_{i \min}}{S}\right)^2 / (S-1)} / \frac{\sum_{i=1}^S d_{i \min}}{S}$ | Nearest equability | This paper |
| Evenness of nearest neighbor distance | $FNEve$<br>[evenNND] | Equality in nearest distances | $\left(\sum_{i=1}^S \min\left(\frac{d_{i \min}}{\sum_{i=1}^{S-1} d_{i \min}}, \frac{1}{S}\right) - \frac{1}{S}\right) / \left(1 - \frac{1}{S}\right)$ | Nearest equability | (Weiher 2011) |

**Table A2.** Metrics that combine species richness with one other basic element.

| Metric | Symbol | Description | Formula | Quantities | Source |
| --- | --- | --- | --- | --- | --- |
| **A. Magnitude** |  |  |  |  |  |
| Total abundance | N | Total number of individuals | $\sum_{i=1}^S n_i$ | Identity, numbers | NA |
| Faith's phylogenetic diversity | $\Sigma(P_T)$<br>[PD] | Total branch length | $\sum_i \sum_{b \in b(S_i)} L_{ij}$ | Identity, total divergence | (Faith 1992) |
| Chao's phylogenetic richness | M(PR) | Effective number of phylogenetically-independent species | $\frac{\sum_i \sum_{b \in b(S_i)} L_{ij}}{T}$ | Identity, total divergence | (Scheiner et al. 2017a) |
| Extensive quadratic entropy | $\Sigma(P_P)$<br>[F] | Sum of pairwise phylogenetic divergences | $\sum_{i=1}^S \sum_{j=1}^S \varphi_{ij}$ | Identity, pairwise divergence | (Izsák and Papp 2000) |
| Phylogenetic species richness | PSR | Sum of unshared branch lengths | $\sum_{i=1}^S \sum_{j=1}^S \psi_{ij}$ | Identity, pairwise divergence | (Helmus et al. 2007) |
| Nearest taxon distance | $\Sigma(P_N)$ | Sum of shortest phylogenetic divergences | $\sum_{i=1}^S \varphi_{i \min}$ | Identity, nearest divergence | This paper |
| Functional attribute diversity | $\Sigma(T_{T/P})$<br>[FAD] | Sum of pairwise trait distances | $\sum_{i=1}^S \sum_{j=1}^S d_{ij}$ | Identity, total or pairwise dispersion | (Walker et al. 1999) |
| Functional richness | FRic | Trait-space convex hull volume | Quickhull algorithm | Identity, total dispersion | (Cornwell et al. 2006) |
| Nearest trait dispersion | $\Sigma(T_N)$ | Sum of shortest trait distances between taxa | $\sum_{i=1}^S d_{i \min}$ | Identity, nearest dispersion | This paper |
| Functional diversity | FD | Sum of the branch lengths of a trait-space minimum spanning tree | $\sum_{l=1}^{S-1} \mu_{l \min}$ | Identity, nearest dispersion | (Petchey and Gaston 2002) |
| **B. Variability** |  |  |  |  |  |
| Hill diversity | ${}^qD(A)$ | Effective number of equally-abundant species | $\left( \sum_{i=1}^S \left( \frac{n_i}{\sum_{i=1}^S n_i} \right)^q \right)^{1/(1-q)}$ | Identity, evenness | (Hill 1973) |
| Phylogenetic Hill diversity | ${}^qD(P_T)$<br>[${}^qD(P)$] | Effective number of equally phylogenetically-distinct species | $\left( \sum_i \left( \frac{\sum_{b \in b(S_i)} L_{ij}}{\sum_i \sum_{b \in b(S_i)} L_{ij}} \right)^q \right)^{1/(1-q)}$ | Identity, total regularity | (Scheiner 2012) |
| Pairwise<br>phylogenetic Hill<br>diversity | ${}^qD(P_P)$ | Effective number of<br>equally phylogenetically-distinct species | $\left(1 + \sqrt{1 + 4 \left( \sum_{i=1}^S \sum_{j=1}^S \left( \frac{\varphi_{ij}}{\sum_{i=1}^S \sum_{j=1}^S \varphi_{ij}} \right)^q \right)^{1/(1-q)}} \right) / 2$ | Identity, pairwise<br>regularity | This paper |
| Nearest taxon Hill<br>diversity | ${}^qD(P_N)$ | Effective number of<br>equally phylogenetically-distinct species | $\left( \sum_{i=1}^S \left( \frac{\varphi_{i \min}}{\sum_{i=1}^S \varphi_{i \min}} \right)^q \right)^{1/(1-q)}$ | Identity, nearest<br>regularity | This paper |
| Functional-trait<br>Hill diversity | ${}^qD(T_T)$<br>[${}^qD(T^*)$] | Effective number of<br>equally functionally-<br>distinct species | $\left( \sum_{i=1}^S \left( \frac{d_i}{\sum_{i=1}^S d_i} \right)^q \right)^{1/(1-q)}$ | Identity, total<br>equability | (Presley et al.<br>2014) |
| Pairwise<br>functional-trait Hill<br>diversity | ${}^qD(T_P)$<br>[${}^qD(T)$] | Effective number of<br>equally functionally-<br>distinct species | $\left(1 + \sqrt{1 + 4 \left( \sum_{i=1}^S \sum_{j=1}^S \left( \frac{d_{ij}}{\sum_{i=1}^S \sum_{j=1}^S d_{ij}} \right)^q \right)^{1/(1-q)}} \right) / 2$ | Identity, pairwise<br>equability | (Scheiner et al.<br>2017b) |
| Nearest functional-trait Hill diversity | ${}^qD(T_N)$<br>[${}^qD(F)$] | Effective number of<br>distinct species | $\left( \sum_{i=1}^S \left( \frac{d_{i \min}}{\sum_{i=1}^S d_{i \min}} \right)^q \right)^{1/(1-q)}$ | Identity, nearest<br>equability | (Scheiner 2012) |

**Table A3.**
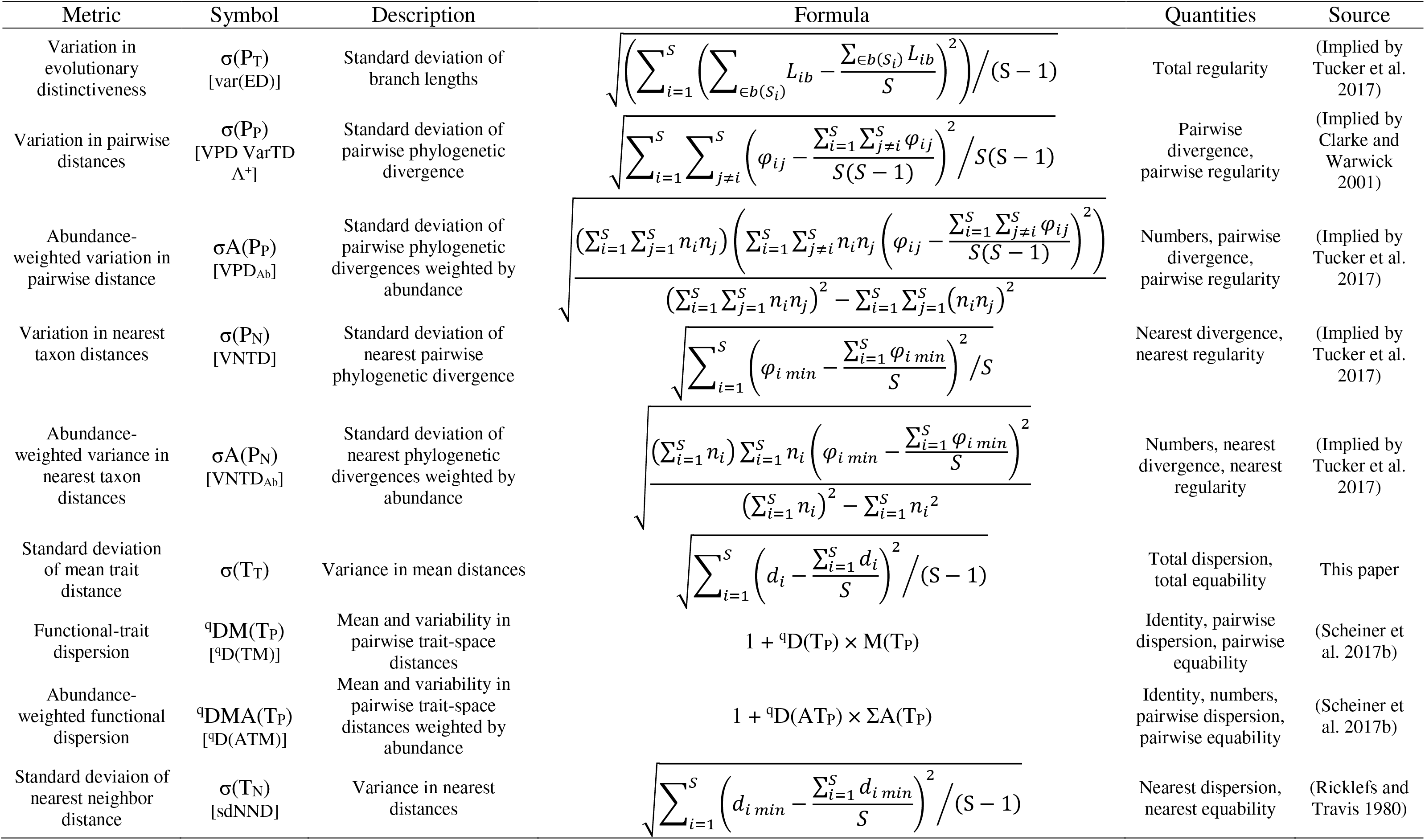
Metrics that combine magnitude and variability of the same data type

| Metric | Symbol | Description | Formula | Quantities | Source |
| --- | --- | --- | --- | --- | --- |
| Variation in evolutionary distinctiveness | $\sigma(P_T)$<br>[var(ED)] | Standard deviation of branch lengths | $\sqrt{\left(\sum_{i=1}^S \left(\sum_{b \in b(S_i)} L_{ib} - \frac{\sum_{b \in b(S_i)} L_{ib}}{S}\right)^2\right) / (S-1)}$ | Total regularity | (Implied by Tucker et al. 2017) |
| Variation in pairwise distances | $\sigma(P_P)$<br>[VPD VarTD $\Lambda^+$] | Standard deviation of pairwise phylogenetic divergence | $\sqrt{\sum_{i=1}^S \sum_{j \neq i}^S \left(\varphi_{ij} - \frac{\sum_{i=1}^S \sum_{j \neq i}^S \varphi_{ij}}{S(S-1)}\right)^2 / S(S-1)}$ | Pairwise divergence, pairwise regularity | (Implied by Clarke and Warwick 2001) |
| Abundance-weighted variation in pairwise distance | $\sigma A(P_P)$<br>[VPD<sub>Ab</sub>] | Standard deviation of pairwise phylogenetic divergences weighted by abundance | $\sqrt{\frac{\left(\sum_{i=1}^S \sum_{j=1}^S n_i n_j\right) \left(\sum_{i=1}^S \sum_{j \neq i}^S n_i n_j \left(\varphi_{ij} - \frac{\sum_{i=1}^S \sum_{j \neq i}^S \varphi_{ij}}{S(S-1)}\right)^2\right)}{\left(\sum_{i=1}^S \sum_{j=1}^S n_i n_j\right)^2 - \sum_{i=1}^S \sum_{j=1}^S (n_i n_j)^2}}$ | Numbers, pairwise divergence, pairwise regularity | (Implied by Tucker et al. 2017) |
| Variation in nearest taxon distances | $\sigma(P_N)$<br>[VNTD] | Standard deviation of nearest pairwise phylogenetic divergence | $\sqrt{\sum_{i=1}^S \left(\varphi_{i \min} - \frac{\sum_{i=1}^S \varphi_{i \min}}{S}\right)^2 / S}$ | Nearest divergence, nearest regularity | (Implied by Tucker et al. 2017) |
| Abundance-weighted variance in nearest taxon distances | $\sigma A(P_N)$<br>[VNTD<sub>Ab</sub>] | Standard deviation of nearest phylogenetic divergences weighted by abundance | $\sqrt{\frac{\left(\sum_{i=1}^S n_i\right) \sum_{i=1}^S n_i \left(\varphi_{i \min} - \frac{\sum_{i=1}^S \varphi_{i \min}}{S}\right)^2}{\left(\sum_{i=1}^S n_i\right)^2 - \sum_{i=1}^S n_i^2}}$ | Numbers, nearest divergence, nearest regularity | (Implied by Tucker et al. 2017) |
| Standard deviation of mean trait distance | $\sigma(T_T)$ | Variance in mean distances | $\sqrt{\sum_{i=1}^S \left(d_i - \frac{\sum_{i=1}^S d_i}{S}\right)^2 / (S-1)}$ | Total dispersion, total equability | This paper |
| Functional-trait dispersion | ${}^qDM(T_P)$<br>[${}^qD(TM)$] | Mean and variability in pairwise trait-space distances | $1 + {}^qD(T_P) \times M(T_P)$ | Identity, pairwise dispersion, pairwise equability | (Scheiner et al. 2017b) |
| Abundance-weighted functional dispersion | ${}^qDMA(T_P)$<br>[${}^qD(ATM)$] | Mean and variability in pairwise trait-space distances weighted by abundance | $1 + {}^qD(AT_P) \times \Sigma A(T_P)$ | Identity, numbers, pairwise dispersion, pairwise equability | (Scheiner et al. 2017b) |
| Standard deviation of nearest neighbor distance | $\sigma(T_N)$<br>[sdNND] | Variance in nearest distances | $\sqrt{\sum_{i=1}^S \left(d_{i \min} - \frac{\sum_{i=1}^S d_{i \min}}{S}\right)^2 / (S-1)}$ | Nearest dispersion, nearest equability | (Ricklefs and Travis 1980) |

**Table A4.**
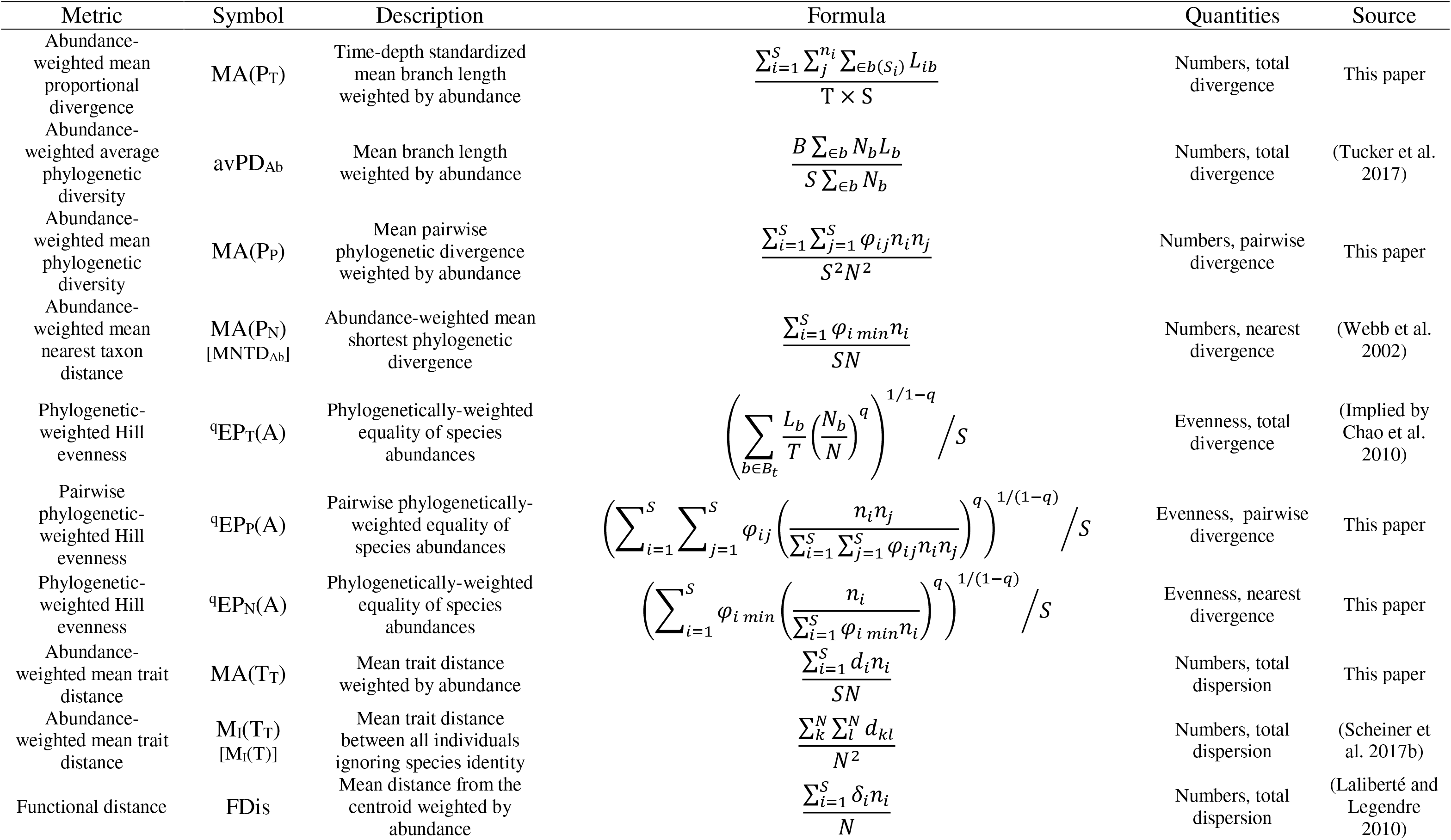
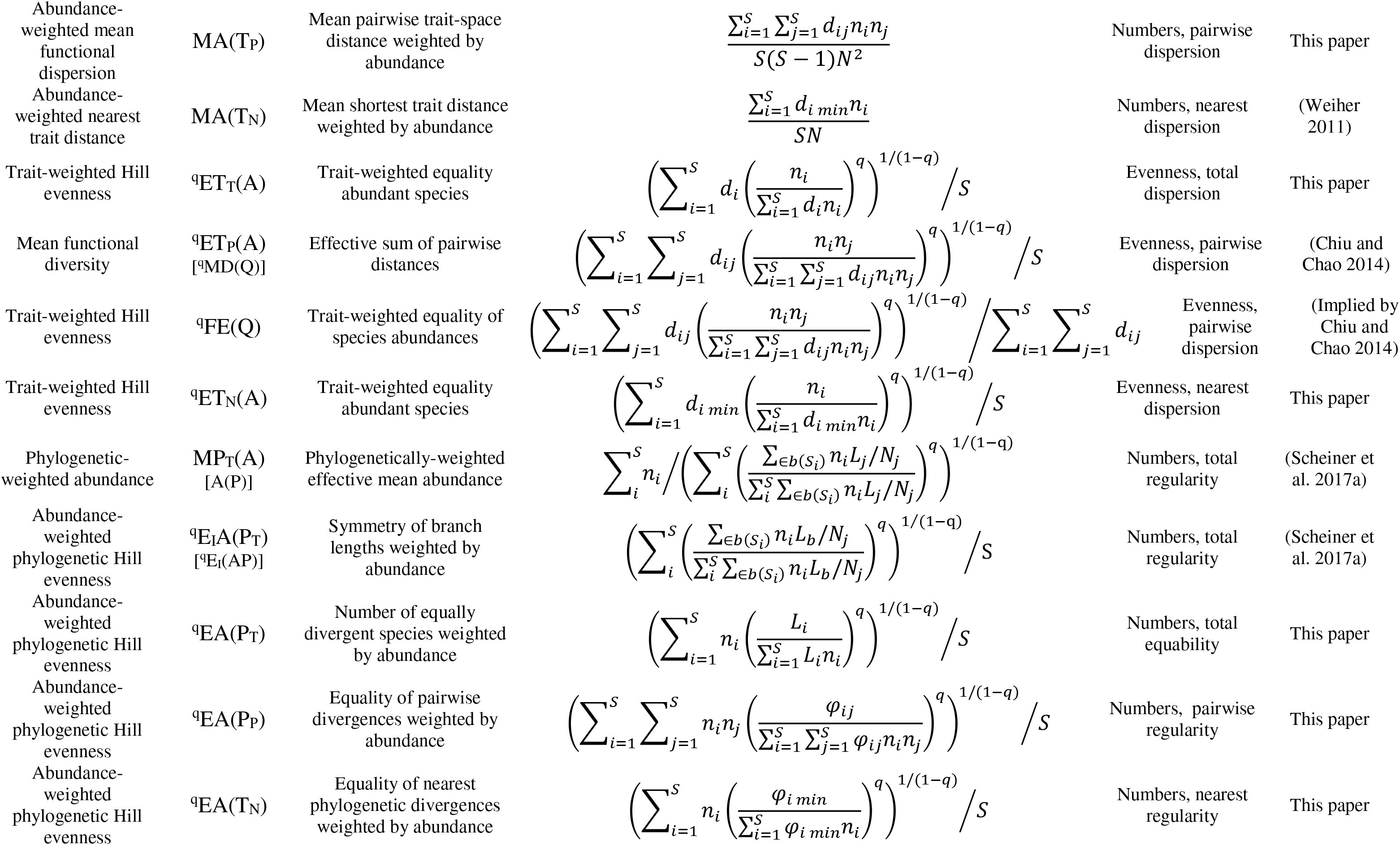
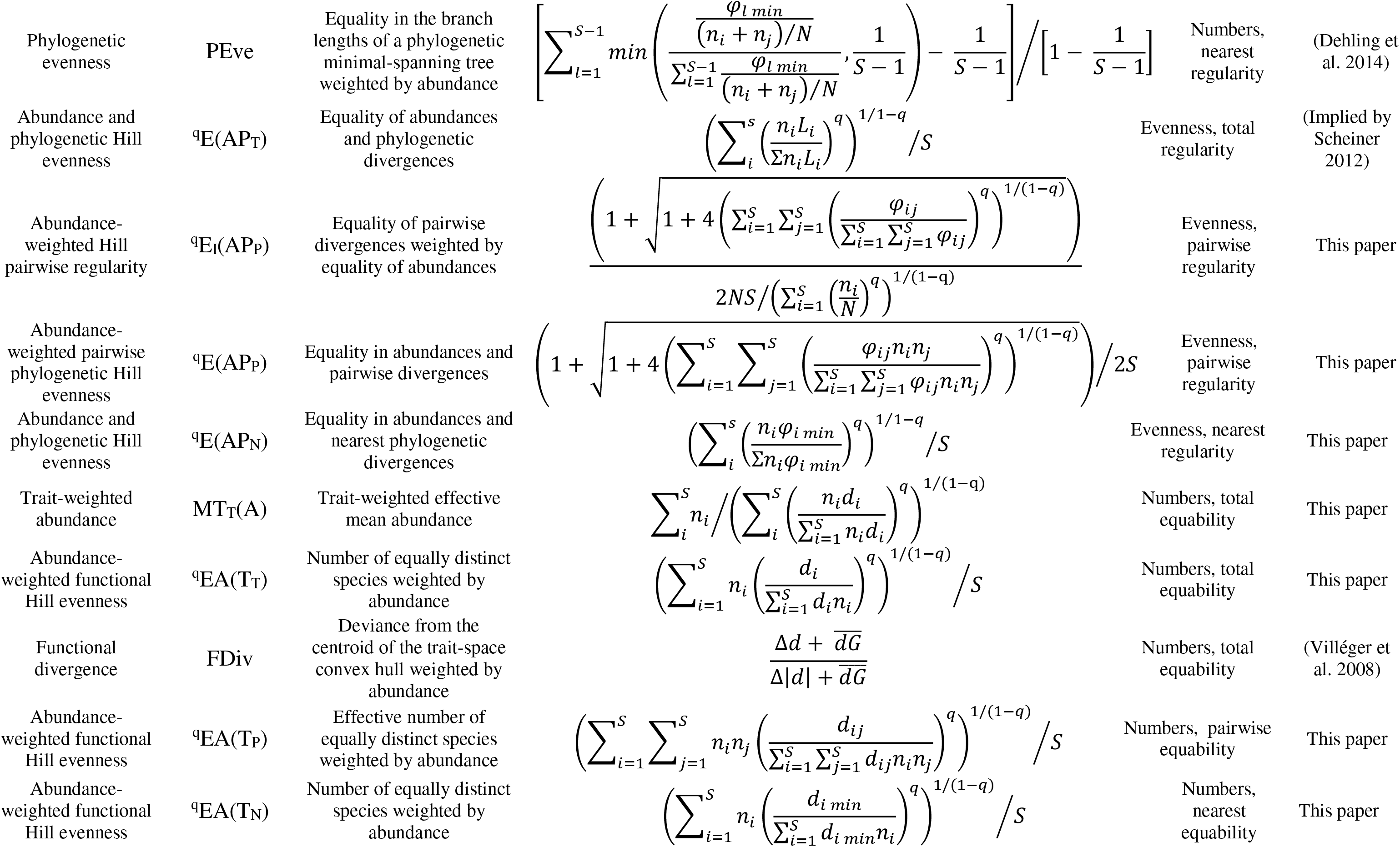
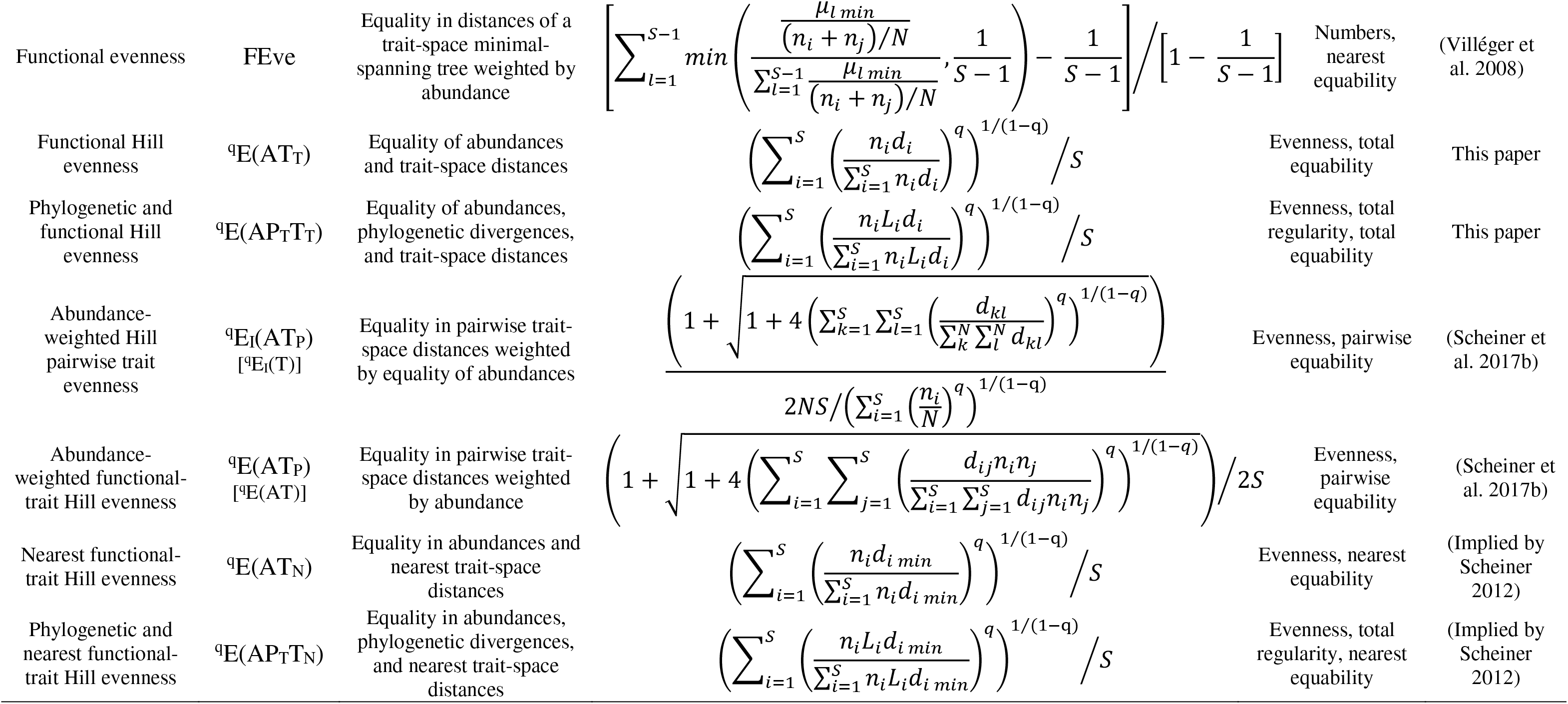
Metrics that combine abundance with one other basic element involving phylogeny or traits.

**Table A5.** Metrics that combine species richness and abundance with one or more other basic elements involving phylogeny or traits.

| Metric | Symbol | Description | Formula | Quantities | Source |
| --- | --- | --- | --- | --- | --- |
| Abundance-weighted phylogenetic diversity | $\Sigma(\text{AP}_T)$<br>[$\Delta\text{nPD}$] | The sum of branch lengths weighted by abundance | $\sum_{b \in B} N_b L_b$ | Identity, numbers, total divergence | (Barker 2002) |
| Abundance-weighted phylogenetic diversity | $\Sigma\text{A}(\text{P}_T)$<br>[$\text{PD}_{\text{Ab}}$] | The sum of branch lengths weighted by abundance | $\frac{B \sum_{b \in B} N_b L_b}{\sum_{b \in B} N_b}$ | Identity, numbers, total divergence | (Vellend et al. 2011) |
| Rao's quadratic phylogenetic entropy | $\Sigma\text{A}(\text{P}_P)$<br>[$\text{MPD}_{\text{Ab}}$] | Expected phylogenetic divergence of two randomly drawn individuals | $\frac{\sum_{i=1}^S \sum_{j=1}^S \varphi_{ij} n_i n_j}{N^2}$ | Identity, numbers, pairwise divergence | (Clarke and Warwick 1998) |
| Phylogenetic species evenness | PSE | Pairwise unshared branch lengths weighted by abundance | $\frac{S \sum_{i=1}^S \sum_{j=1}^S \psi_{ij} n_i n_j}{(S-1)N^2}$ | Identity, numbers, pairwise divergence | (Helmus et al. 2007) |
| Abundance-weighted nearest taxon distance | $\Sigma\text{A}(\text{P}_N)$ | Abundance-weighted sum of the shortest phylogenetic divergences | $\frac{\sum_{i=1}^S \varphi_{i \min} n_i}{N}$ | Numbers nearest divergence | (Implied by Webb et al. 2002) |
| Phylogenetic-weighted Hill diversity | ${}^q\text{DP}_T(\text{A})$<br>[${}^q\bar{D}(T)$] | Phylogenetic-weighted effective number of equally abundant species | $\left( \sum_{b \in B_t} \frac{L_b}{T} \left( \frac{N_b}{N} \right)^q \right)^{1/(1-q)}$ | Identity, evenness, total divergence | (Chao et al. 2010) |
| Pairwise phylogenetic-weighted Hill diversity | ${}^q\text{DP}_P(\text{A})$ | Effective number of equally abundant species weighted by pairwise phylogenetic divergences | $\left( \sum_{i=1}^S \sum_{j=1}^S \varphi_{ij} \left( \frac{n_i n_j}{\sum_{i=1}^S \sum_{j=1}^S \varphi_{ij} n_i n_j} \right)^q \right)^{1/(1-q)}$ | Identity, evenness, pairwise divergence | This paper |
| Phylogenetic-weighted Hill diversity | ${}^q\text{DP}_N(\text{A})$ | Effective number of equally abundant species weighted by nearest phylogenetic divergences | $\left( \sum_{i=1}^S \varphi_{i \min} \left( \frac{n_i}{\sum_{i=1}^S \varphi_{i \min} n_i} \right)^q \right)^{1/(1-q)}$ | Identity, evenness, nearest dispersion | This paper |
| Abundance-weighted trait dispersion | $\Sigma\text{A}(\text{T}_T)$ | The sum of mean species distances weighted by abundance | $\frac{\sum_{i=1}^S d_i n_i}{N}$ | Identity, numbers, total dispersion | This paper |
| Rao's quadratic trait entropy | $\Sigma\text{A}(\text{T}_P)$<br>[$\text{Q}(\text{T})$] | Mean trait distance between any two randomly drawn individuals | $\frac{\sum_{i=1}^S \sum_{j=1}^S d_{ij} n_i n_j}{N^2}$ | Identity, numbers, pairwise dispersion | (Chiu and Chao 2014) |
| Abundance-weighted nearest trait distance | $\Sigma A(T_N)$ | Mean shortest trait distance weighted by abundance | $\frac{\sum_{i=1}^S d_{i \min} n_i}{N}$ | Identity, numbers, nearest dispersion | (Implied by Weiher 2011) |
| Trait-weighted Hill diversity | ${}^qDT_T(A)$ | Trait-weighted effective number of equally abundant species | $\left( \sum_{i=1}^S d_i \left( \frac{n_i}{\sum_{i=1}^S d_i n_i} \right)^q \right)^{1/(1-q)}$ | Identity, evenness, total dispersion | This paper |
| Trait-weighted Hill diversity | ${}^qDT_P(A)$<br>[${}^qFD(Q)$] | Trait-weighted effective number of equally abundant species | $\left( \sum_{i=1}^S \sum_{j=1}^S d_{ij} \left( \frac{n_i n_j}{\sum_{i=1}^S \sum_{j=1}^S d_{ij} n_i n_j} \right)^q \right)^{1/(1-q)}$ | Identity, evenness, pairwise dispersion | (Chiu and Chao 2014) |
| Trait-weighted Hill diversity | ${}^qDT_N(A)$ | Trait-weighted effective number of equally abundant species | $\left( \sum_{i=1}^S d_{i \min} \left( \frac{n_i}{\sum_{i=1}^S d_{i \min} n_i} \right)^q \right)^{1/(1-q)}$ | Identity, evenness, nearest dispersion | This paper |
| Abundance-weighted phylogenetic Hill diversity | ${}^qD_I A(P_T)$<br>[${}^qD_I(AP)$] | Effective number of phylogenetically-distinct species weighted by abundance | $\left( \sum_i \left( \frac{\sum_{b \in b(S_i)} n_i L_b / N_b}{\sum_i \sum_{b \in b(S_i)} n_i L_b / N_b} \right)^q \right)^{1/(1-q)}$ | Identity, numbers, total regularity | (Scheiner 2012) |
| Abundance-weighted phylogenetic Hill diversity | ${}^qDA(P_T)$ | Effective number of equally divergent species weighted by abundance | $\left( \sum_{i=1}^S n_i \left( \frac{L_i}{\sum_{i=1}^S L_i n_i} \right)^q \right)^{1/(1-q)}$ | Identity, numbers, total regularity | This paper |
| Abundance-weighted phylogenetic Hill diversity | ${}^qDA(P_P)$ | Effective number of equally divergent species weighted by abundance | $\left( \sum_{i=1}^S \sum_{j=1}^S n_i n_j \left( \frac{\varphi_{ij}}{\sum_{i=1}^S \sum_{j=1}^S \varphi_{ij} n_i n_j} \right)^q \right)^{1/(1-q)}$ | Identity, numbers, pairwise regularity | This paper |
| Abundance-weighted functional Hill diversity | ${}^qDA(P_N)$ | Effective number of equally distinct species weighted by abundance | $\left( \sum_{i=1}^S n_i \left( \frac{\varphi_{i \min}}{\sum_{i=1}^S \varphi_{i \min} n_i} \right)^q \right)^{1/(1-q)}$ | Identity, numbers, nearest regularity | This paper |
| Abundance and phylogenetic Hill diversity | ${}^qD(AP_T)$<br>[${}^qD(AP)$] | Effective number of equally abundant and phylogenetically distinct species | $\left( \sum_i \left( \frac{n_i L_i}{\sum n_i L_i} \right)^q \right)^{1/1-q}$ | Identity, evenness, total regularity | (Scheiner 2012) |
| Abundance-weighted phylogenetic Hill diversity | ${}^qD_I(AP_P)$ | Effective number of phylogenetically-distinct species weighted by the effective abundance | $\frac{\left( 1 + \sqrt{1 + 4 \left( \sum_{i=1}^S \sum_{j=1}^S \left( \frac{\varphi_{ij}}{\sum_{i=1}^S \sum_{j=1}^S \varphi_{ij}} \right)^q} \right)^{1/(1-q)}} \right)}{2N / \left( \sum_{i=1}^S \left( \frac{n_i}{N} \right)^q \right)^{1/(1-q)}}$ | Identity, numbers, pairwise regularity | This paper |
| Abundance-weighted pairwise phylogenetic Hill diversity | ${}^qD(AP_P)$ | Effective number of equally abundant and phylogenetically distinct species | $\left(1 + \sqrt{1 + 4 \left( \sum_{i=1}^S \sum_{j=1}^S \left( \frac{\varphi_{ij} n_i n_j}{\sum_{i=1}^S \sum_{j=1}^S \varphi_{ij} n_i n_j} \right)^q \right)^{1/(1-q)}} \right) / 2$ | Identity, evenness, pairwise regularity | This paper |
| Abundance and nearest taxon Hill diversity | ${}^qD(AP_N)$ | Effective number of equally abundant and phylogenetically distinct species | $\left( \sum_{i=1}^S \left( \frac{n_i \varphi_{i \min}}{\sum_{i=1}^S n_i \varphi_{i \min}} \right)^q \right)^{1/(1-q)}$ | Identity, evenness, nearest regularity | This paper |
| Abundance-weighted functional Hill diversity | ${}^qDA(T_T)$ | Effective number of equally distinct species weighted by abundance | $\left( \sum_{i=1}^S n_i \left( \frac{d_i}{\sum_{i=1}^S d_i n_i} \right)^q \right)^{1/(1-q)}$ | Identity, numbers, total equability | This paper |
| Abundance-weighted functional Hill diversity | ${}^qDA(T_P)$ | Effective number of equally distinct species weighted by abundance | $\left( \sum_{i=1}^S \sum_{j=1}^S n_i n_j \left( \frac{d_{ij}}{\sum_{i=1}^S \sum_{j=1}^S d_{ij} n_i n_j} \right)^q \right)^{1/(1-q)}$ | Identity, numbers, pairwise equability | This paper |
| Abundance-weighted functional Hill diversity | ${}^qDA(T_N)$ | Number of equally distinct species weighted by abundance | $\left( \sum_{i=1}^S n_i \left( \frac{d_{i \min}}{\sum_{i=1}^S d_{i \min} n_i} \right)^q \right)^{1/(1-q)}$ | Identity, numbers, nearest equability | This paper |
| Abundance and functional Hill diversity | ${}^qD(AT_T)$ | Effective number of equally abundant and functionally distinct species | $\left( \sum_{i=1}^S \left( \frac{n_i d_i}{\sum_{i=1}^S n_i d_i} \right)^q \right)^{1/(1-q)}$ | Identity, evenness, total equability | This paper |
| Abundance, phylogenetic and functional Hill diversity | ${}^qD(AP_T T_T)$ | Effective number of equally abundant, phylogenetically distinct and functionally distinct species | $\left( \sum_{i=1}^S \left( \frac{n_i L_i d_i}{\sum_{i=1}^S n_i L_i d_i} \right)^q \right)^{1/(1-q)}$ | Identity, evenness, total regularity, total equability | This paper |
| Abundance-weighted Hill pairwise trait diversity | ${}^qD_I(AT_P)$<br>[${}^qD_I(T)$] | Effective number of functionally distinct species weighted by the effective abundance | $\frac{\left(1 + \sqrt{1 + 4 \left( \sum_{k=1}^S \sum_{l=1}^S \left( \frac{d_{kl}}{\sum_{k=1}^S \sum_{l=1}^S d_{kl}} \right)^q \right)^{1/(1-q)}} \right)}{2N / \left( \sum_{i=1}^S \left( \frac{n_i}{N} \right)^q \right)^{1/(1-q)}}$ | Identity, evenness, pairwise equability | (Scheiner et al. 2017b) |
| Abundance-weighted functional Hill diversity | ${}^qD(AT_P)$<br>[${}^qD(AT)$] | Variability in pairwise distances weighted by abundance | $\left(1 + \sqrt{1 + 4 \left( \sum_{i=1}^S \sum_{j=1}^S \left( \frac{d_{ij} n_i n_j}{\sum_{i=1}^S \sum_{j=1}^S d_{ij} n_i n_j} \right)^q \right)^{1/(1-q)}} \right) / 2$ | Identity, numbers, pairwise equability | (Scheiner et al. 2017b) |
| Abundance and<br>nearest functional<br>Hill diversity | ${}^qD(AT_N)$ | Effective number of<br>equally abundant and<br>functionally distinct<br>species | $\left( \sum_{i=1}^S \left( \frac{n_i d_{i \min}}{\sum_{i=1}^S n_i d_{i \min}} \right)^q \right)^{1/(1-q)}$ | Identity, evenness,<br>nearest equability | (Scheiner 2012) |
| Abundance,<br>phylogenetic and<br>nearest functional<br>Hill diversity | ${}^qD(AP^T T_N)$<br>[${}^qD(APF)$] | Effective number of<br>equally abundant,<br>phylogenetically distinct,<br>and functionally distinct<br>species | $\left( \sum_{i=1}^S \left( \frac{n_i L_i d_{i \min}}{\sum_{i=1}^S n_i L_i d_{i \min}} \right)^q \right)^{1/(1-q)}$ | Identity, evenness,<br>total regularity,<br>nearest equability | (Scheiner 2012) |

